# Image Analysis for Non-Neoplastic Kidney Disease: Utilizing Morphological Segmentation to Improve Quantification of Interstitial Fibrosis

**DOI:** 10.1101/2025.11.10.686700

**Authors:** Nazanin Mola, Maya Maya Barbosa Silva, Sabine Leh, Hrafn Weishaupt

## Abstract

Interstitial fibrosis (IF) is a hallmark of chronic kidney disease (CKD) and a strong predictor of progression to end-stage kidney disease (ESKD). Current biopsy-based IF assessments rely on subjective visual estimations, limiting reproducibility. Sirius Red staining is widely used for visualizing fibrotic tissue, yet its application in digital pathology is limited by non-specific staining. This study investigates the impact of cortical structure segmentation on fibrosis quantification in Sirius Red-stained, non-neoplastic kidney biopsies. Fibrosis measurements before and after segmentation were compared using two image analysis methods (stain deconvolution and red-green), with ground truth fibrosis measured by point counting and expert pathologist grading.

Excluding non-interstitial structures led to a significant reduction of quantified fibrosis and improved correlation with pathology grading and point counting for both the stain deconvolution and the red green method. Bland-Altman analysis showed reduced bias after segmentation: for the deconvolution method, mean difference decreased from +2.5% (95% LoA: -8% to +13%) to +1% (-7% to +9%); for the red-green method, from +3% (-10% to +16%) to +1% (-8% to +10%). Correlation with pathology grading also improved (Spearmans ρ rose from 0.55 to 0.58 for deconvolution and from 0.53 to 0.59 for red-green).

These findings confirm that targeted segmentation enhances the accuracy and consistency of automated fibrosis assessment, supporting its integration into digital pathology workflows as a critical step toward reliable quantification of fibrosis in kidney disease.

## 2. Introduction

All chronic kidney diseases develop over time an accumulation of connective tissue in the interstitium, known as interstitial fibrosis (IF). The connection between interstitial fibrosis and kidney failure has long been recognized^1,2,3^, with IF grading being an important prognostic marker for the development of end-stage kidney disease (ESKD) in various non-neoplastic kidney disorders and transplant biopsies^4-6^. Consequently, an accurate measurement of a biopsy’s IF level is essential for both patient management and clinical research^7, 8-14^.

However, current biopsy scoring systems categorize the extent of cortical interstitial fibrosis based on a visual estimation of the percentage of affected tissue, usually into four categories: minimal (≤5%), mild (6–25%), moderate (26–50%), and severe (>50%)^15,16^. Commonly referred to as “eyeballing”, these visual methods, while rapid and straightforward, are subjected to high inter-observer variability and limited reproducibility^17,18,19^, which undermines their utility as a quantitative digital biomarker. Additionally, there seems to be a lack of standardization on how to effectively quantify said percentage of affected tissue as discussed and illustrated by Farris, et al.^20^ in their two different visual estimation approaches.

To address these limitations, various image analysis and machine learning (ML)^21-24^ approaches have been explored in different histological settings. Among the histological stains commonly used for the visualization and computer-aided quantification of fibrotic tissue, Picrosirius Red (hereafter ‘Sirius Red’) stands out as a robust and easy-to-use choice^19^, with research indicating it as one of the best stains in the correlation to both clinical parameters and eyeballing results^25^. Its high specificity to collagen fibre types I and III enables a clear/contrastive visualization of collagenous rich areas of the tissue section, particularly when viewed under polarized light microscopy. Routine whole-slide image (WSI) scanners, however, generally do not yet support polarized light imaging and, therefore, digitization of polarized microscopy sections remains impractical. Nonetheless, Sirius Red retains its high correlation to expert evaluations and clinical biomarker levels in chronic kidney disease (CKD)^25^, albeit also more frequently staining non-interstitial structures, specifically tubules’ epithelial cell nuclei and glomerular structures^26,27^.

With respect to unspecific staining, a few studies have attempted to exclude the largest structures representing non-specific staining (glomeruli) and, in addition, non-relevant fibrotic structures (perivascular fibrotic cuffs), in order to better approximate the true region of interest for fibrosis quantification, namely the kidney interstitium^28-30^. Nevertheless, to the best of our knowledge, the effectiveness of structural segmentation in improving automatic fibrosis measurement has not been systematically investigated, and existing studies often fall short by not performing a full segmentation of all major kidney structures nor comparing fibrosis quantification results before and after segmentation to explicitly demonstrate the impact of this step on measurement accuracy.

To address this gap, the current study sought to specifically evaluate the effect of cortical structure segmentation prior to fibrosis quantification, utilizing unpolarized Sirius Red-stained WSIs, and testing the performance of fibrosis quantification with and without the removal of non-interstitial structures with respect to (i) two conventional image-analysis methods, (ii) two pathologist-defined fibrosis measurements, and (iii) serum creatinine. Together, we believe that these experiments present a more systematic insight into the benefits of structure segmentation and towards objective and interstitium-specific strategies of fibrosis quantification in Sirius Red-stained non-neoplastic kidney biopsies.

## 3. Material and methods

To extensively investigate the role of a complete structural segmentation in automated fibrosis quantification methods, we proposed to study its effect across two different image analysis-pipelines, namely stain deconvolution and a red–green method, and with respect to two pathologist-derived reference standards and a relevant clinical variable. Specifically, we proposed to segment (i) glomeruli, (ii) tubules and (iii) big vasculatures, to generate an interstitium-specific quantification framework for non-neoplastic kidney biopsies. To our knowledge, this is the most comprehensive study focused on directly quantifying the impact of a robust non-interstitial structural segmentation step on different Digital image analysis based IF quantification methods.

### 3.1. Dataset

The study included 16 non-neoplastic kidney biopsies obtained from routine diagnostic procedures at the Haukeland University Hospital between 2019 and 2020. An overview of the sample characteristics is provided in Table 1. All samples were native kidney needle biopsies, fixed in formalin, sectioned at 3 μm thickness, and manually stained with Sirius Red following an existing protocol previously published by Farris et al^25^.

**Table 1.** showing clinical data and morphologic characteristics for the 16 kidney biopsies.

| Characteristic | Value |
| --- | --- |
| Serum Creatinine ($\mu\text{mol/L}$) | $306 \pm 275.98$ (range: 75 – 1000, n = 16) |
| Scoring | Minimal: 4, Mild: 4, Moderate: 4, Severe: 4 |
| Cortical area | $5.87 \pm 2.76 \text{ mm}^2$ (mean ±SD) |

The stained tissue sections were digitized using the ScanScope® XT scanner (Aperio Technologies, Inc., Vista, CA, USA) at 40x magnification (0.228 μm/pixel). Slides containing visible artifacts, such as clefts, were identified through visual inspection by a nephropathologist (SL) during the initial sample review and excluded during the case selection process. Case selection aimed for a balanced representation across the spectrum of interstitial fibrosis grades, based on prior Banff grading in diagnostic reports.

### 3.2 Software

Manual annotations were performed in QuPath (v0.5.1), using the wand and brush tools to segment key histological structures. Whole-slide image analyses, including the deconvolution and the red-green methods, were implemented using custom Python scripts using the OpenSlide, OpenCV, Scikit-Image, Histomicstk, Tiffifile and PyVips libraries. Figures were generated with MatPlotLib and/or SeaBorn.

### 3.3 Ground truth (point counting)

A ground truth for interstitial fibrosis was established via point counting^31^ using ImageJ (v1.50i) (National Institutes of Health, Bethesda, MD, USA), by an experienced nephropathologist (SL) manually assigning points within the cortical area to three predefined tissue classes: (i) interstitial matrix (red stained connective tissue); (ii) interstitial space (unstained areas including inflammatory cells and peritubular capillaries); and (iii) others (including glomeruli, and tubules).

In practice, a grid was superimposed over the whole-slide Sirius Red-stained kidney biopsy, with usually 400-500 grid points inside tissue boundaries. At each grid intersection, the tissue located at the upper-right corner (see Supplementary Figure 1 and Table 1) was assigned to one of the aforementioned classes. Finally, interstitial fibrosis was then quantified by the proportion of interstitial matrix (connective tissue) relative to the total number of grid intersections within the cortex.

### 3.4 Analysis

#### 3.4.1 Pre-processing

Each WSI was manually reviewed and non-informative regions, such as the background, glomeruli, tubules, non-cortex tissue, and large blood vessels, were segmented (Figure 1). Both healthy and diseased glomeruli were annotated as a single class. Annotation of cortical tubules was facilitated by a semi-automatic approach where predictions from a preliminary in-house segmentation model trained on publicly available data (KPMP: https://www.kpmp.org/) were used as a starting point to be consequently manually reviewed and corrected.

**Figure 1.**
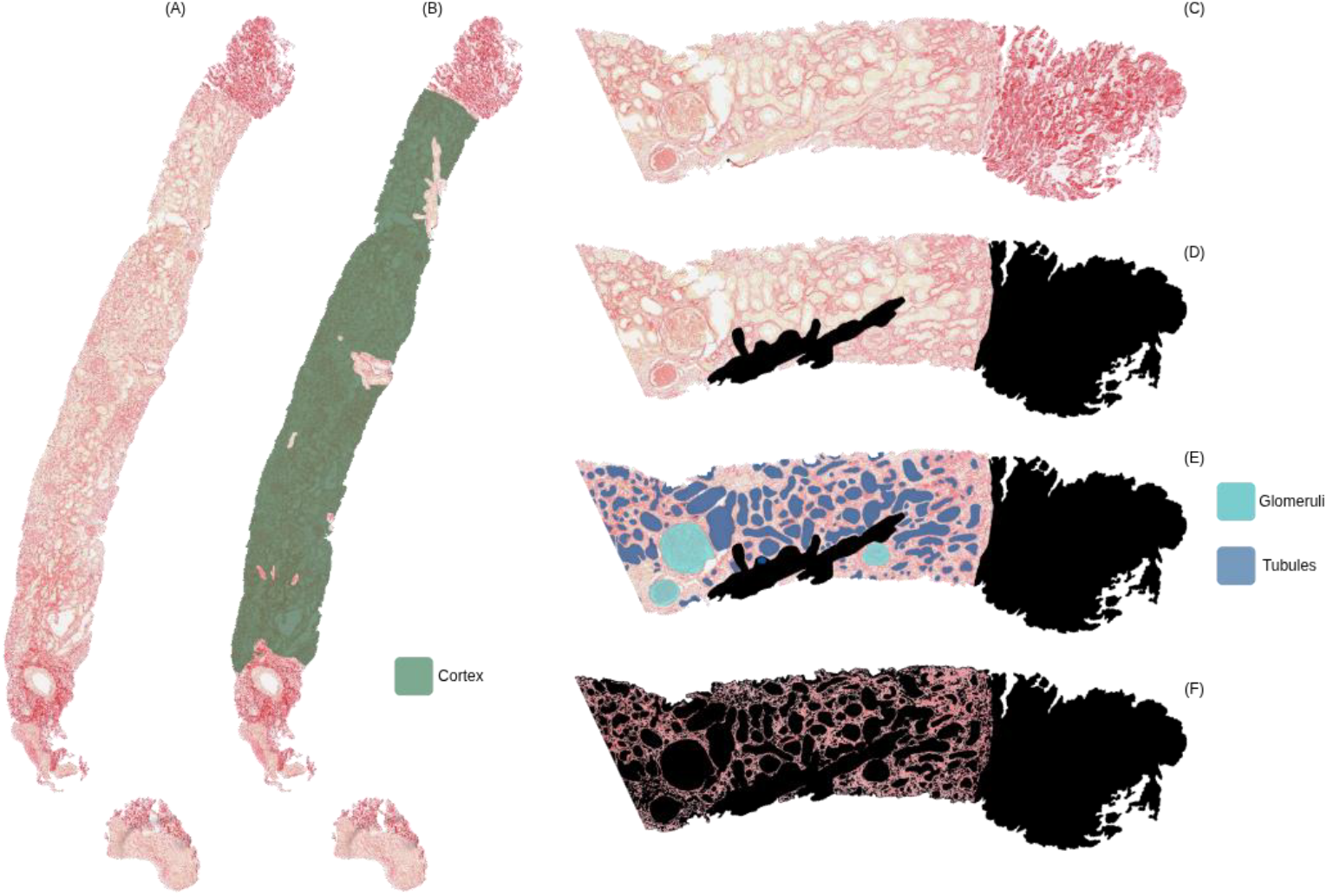
Illustration of the proposed methodology showing: (A) The original needle biopsy stained with Sirius Red; (B) Cortex area annotation; (C) Example ROI from the biopsy; (D) Selection of the valid cortex area, excluding big vessels; (E) Segmentation results including (green) glomeruli and (blue) tubules; and (F) Result of the proposed methodology isolating only the fibrotic tissue.

Following segmentation, whole slide images were tiled into 1024x1024 pixel image patches with a 256-pixel overlap at 40x resolution using the *OpenSlide-Python* (https://github.com/openslide/openslide-python) library. Tiles containing less than 25% cortex were excluded from further analysis, yielding, on average, 121 tiles per biopsy.

#### 3.4.2 Stain Deconvolution method

Stain deconvolution is a computational method for separating individual color components in histologically stained images, isolating each stain’s contribution by unmixing overlapping color channels to enhance the analysis of specific tissue features. (Supplementary Figure 2A-C).

In the current study, the HistomicsTK^32^ library (Kitware Inc.,Clifton Park, NY), was used for stain deconvolution^33^ algorithms, which applies a matrix-based color deconvolution approach to unmix histological stains. The matrix is a stain-to-color map, derived from predefined stains, more specifically, H&E and an additional ‘null’ stain vector for images containing only two stains. This method produces grayscale images, with pixel intensities representing the relative concentration of a given stain, which were consequently thresholded using Otsu’s automatic method to generate a preliminary binary fibrosis mask for downstream pixel-based IF quantification (Figure 2).

**Figure 2.**
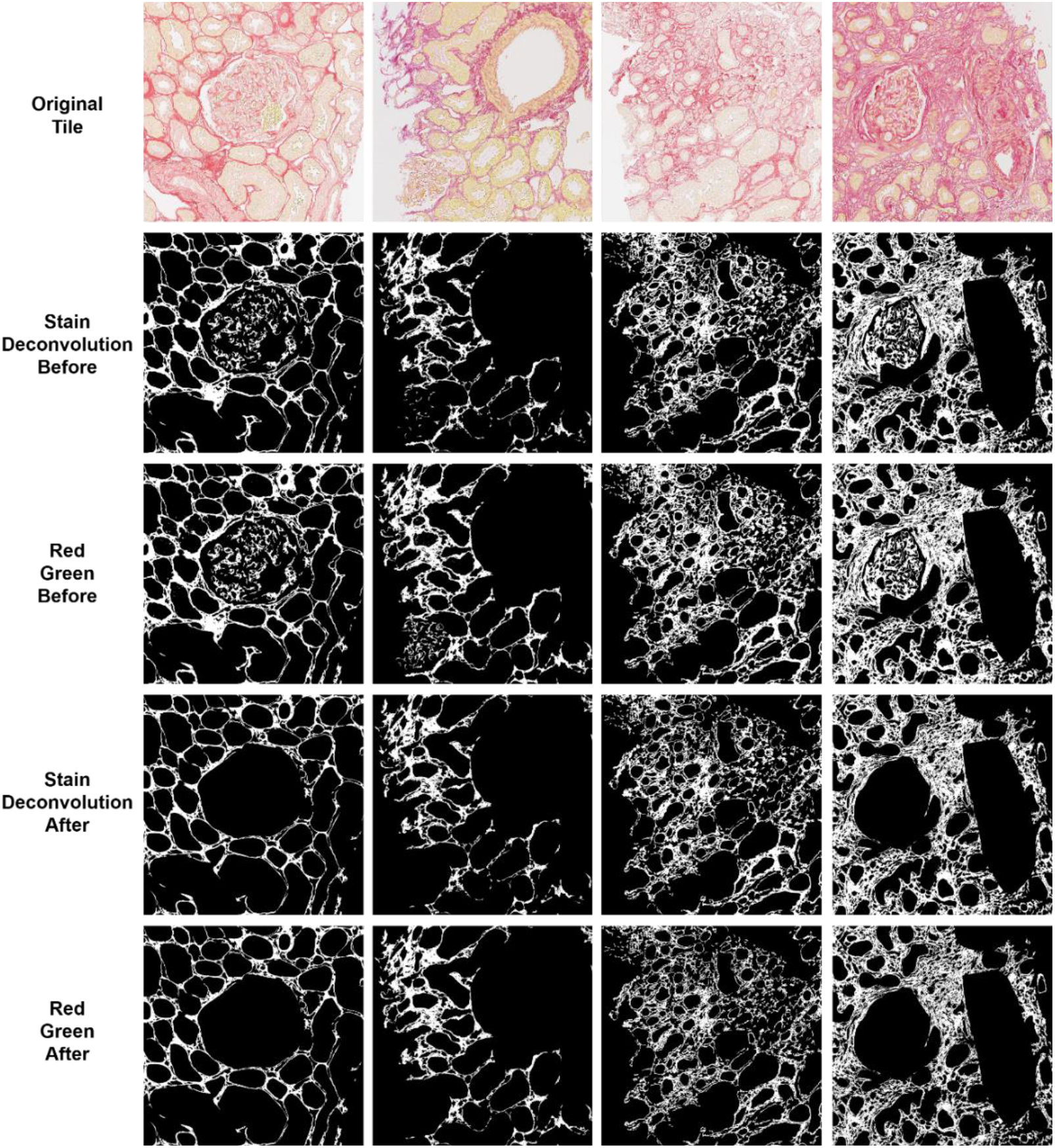
Visualization of two image analysis results. Showing original unprocessed tiles (top row) together with corresponding results produced by the stain deconvolution and red-green method, either before (rows 2 and 3) or after (rows 4 and 5) non-interstitial structure removal.

#### 3.4.3 Red-Green method

Based on the algorithm described by Courtoy, et al.^29^, a “red-green” filtering was used to better contrast the bright positive signal wherever Sirius Red-stained tissue dominates, as most of the background structures (typically stained on a yellow hue due to the Picric acid counter-stain) would have their “R-G” values cluster around zero. Briefly, each image tile had its RGB vectors split into individual channels, using a 32-bit floating point so that the subsequent operations would not clip its dynamic range, and the difference between the red and green channel calculated (Supplementary Figure 2D). Considering the variations in absolute stain intensity, the resultant mask was further “contrast-stretched” using the lower 0.175% and upper 0.175% intensities (0.35% total) as histogram stretching endpoints in a similar fashion to ImageJ’s “Enhance contrast” script. Finally, the resulting image was then automatically thresholded using Otsu’s method and subsequently post-processed to remove any loose components smaller than 100 pixels (Figure 2).

#### 3.4.4. Post-processing

Each of the two image analysis pipelines produced an initial set of tile-level fibrosis masks, from which two result sets were derived for each of the methods: : (i) by directly reassembling the tile-level masks into full-slide binary masks; and (ii) by refining the tile-level masks by excluding both glomeruli and tubules annotated during the pre-processing phase prior to its reassembling into full-slide masks. To quantify the extent of interstitial fibrosis, the proportion of Sirius Red–positive pixels was calculated relative to the cortical area, both with and without structural segmentation:

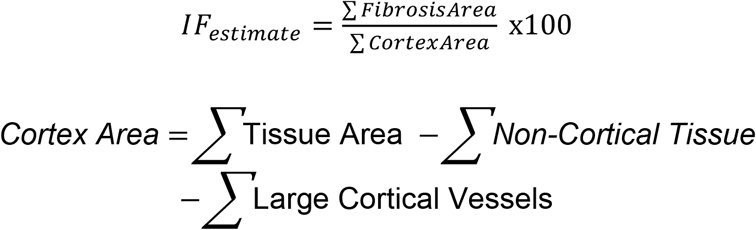

### 3.5. Statistics

Data analysis was performed using descriptive and correlation-based statistical methods. A paired Student’s t-test was used to assess the difference between the two DIA methods. Correlation to the ground-truth was calculated using Spearman’s rank (ρ) and linear regression lines. A one-way analysis of variance (ANOVA) was performed to determine whether there was statistically significant difference, corresponding to a 95% confidence interval, in fibrosis measurements across groups defined by original “eyeballing” pathology grading. Post hoc pairwise comparison was conducted using Tukey’s Honestly Significant Difference (HSD) test to measure the significance of individual group differences. A ρ-value of less than 0.05 was considered statistically significant. To evaluate the agreement between the two computational fibrosis methods and the point-counting-based ground-truth, Bland-Altman analyses were performed. Specifically, these analyses included calculating the mean difference and 95% limits of agreement (mean ± 1.96 x standard deviation), providing insight into both bias and variability across measurements. All statistical analyses and visualizations were conducted using relevant Python libraries, particularly NumPy, SciPy and MatPlotLib.

## 4. Results

To quantify interstitial fibrosis in non-neoplastic Sirius Red-stained kidney biopsies, the current study applied two image analysis methods: a stain deconvolution method, which isolates fibrotic regions based on their optical density, and a red-green method, which enhances contrast for collagen detection and does not rely on predefined stain vectors. Both approaches underwent the same pre-processing steps, including tiling, manual removal of non-cortical tissue and large blood vessels. Finally, fibrosis quantification was performed by automatic thresholding, and final fibrosis percentages were calculated either including or excluding glomeruli and tubules.

### 4.1. Comparison between the methods and their alignment with pathology grading

Both methods, stain deconvolution and the red-green, produced similar trends across the 16 cases, although the red-green method consistently yielded slightly higher fibrosis values than the deconvolution method independent of structural removal, as shown in Figure 3A. As expected, excluding non-interstitial structures resulted in significantly lower fibrosis estimates for each method, for the deconvolution (paired t-test, before vs. after exclusion, p = 2.98 x 10^− 8^) and the red-green method (paired t-test, before vs. after exclusion, p = 9.96 x 10^− 10^) (Figure 3B).

**Figure 3.**
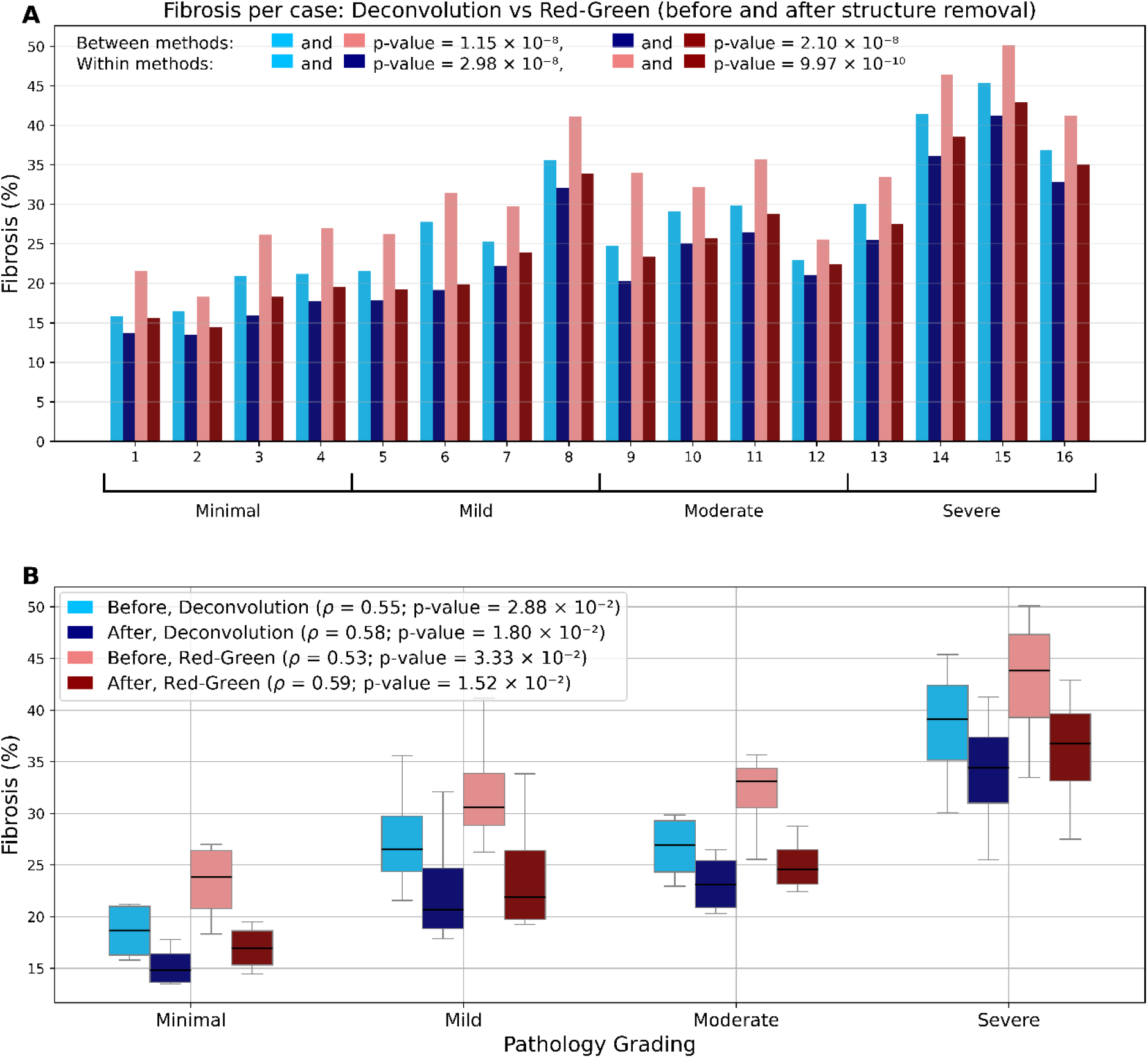
Comparison of fibrosis quantification methods before and after structural segmentation, and correlation with pathology grading. (A) Bar plots showing fibrosis percentages obtained by the deconvolution (blue/purple) and the red-green (pink/red) methods across 16 samples, before and after removal of non-interstitial structures (glomeruli and tubules). Samples are ordered by pathology grade, and within each grade, by ground truth fibrosis percentage. The red-green method consistently yielded higher values than the deconvolution method. Paired t-tests indicate significant differences between the two methods both before (p-value = 1.15 × 10^−8^) and after (p-value = 2.10 × 10^−10^) structural segmentation. Within each method, paired t-tests comparing measurements before and after structural segmentation showed significant, with p-value =2.98 × 10-8 for the deconvolution method and p-value = 9.97 × 10^-10^ for the red-green method. (B) Box plots of fibrosis percentages stratified by pathologist-assigned fibrosis grades (Minimal, Mild, Moderate, Severe), comparing methods before and after structure removal. Spearman’s correlation coefficients (ρ) and p-values show moderate correlation between digital measurements and pathology grading, with slight improvements after structure exclusion.

Another paired t-test comparing the two methods showed a statistically significant difference both before (p = 1.15 x 10^− 8^) and after (p = 2.10 x 10^− 8^) structure removal, suggesting improved agreement following structure exclusion (Figure 3A-B).

Fibrosis estimates from both the deconvolution and the red-green methods increased with higher pathology grades (Figure 3B), as expected. The deconvolution method demonstrated a strong correlation to said grades prior to structure removal which improved slightly afterward, with Spearman’s ρ = 0.55 (p = 2.88 x 10^-2^) increasing to ρ = 0.58 (p =1.80 x 10^-2^). In contrast, the red-green method showed a slightly bigger improvement, with Spearman’s ρ = 0.53 (p = 3.33 x 10^-2^) increasing to 0.59 (p = 1.52 x 10^-2^) after structure removal. ANOVA revealed significant differences in fibrosis estimates from the deconvolution method across pathology grades (F = 9.64, p = 1.62 x 10^−3^). Tukey’s test showed higher values in Severe cases compared to Minimal (p = 9 x 10^−4^), Mild (p = 3.55 x 10^-2^), and Moderate (p = 4.32 x 10^-2^), with no significant differences among the lower grades. The red-green method showed similar patterns with significant fibrosis differences across pathology grades (F = 9.73, p = 1.55 x 10^−3^). Tukey’s test indicated significantly higher fibrosis in Severe cases compared to Minimal (p = 9 x 10^−4^), Mild (p = 2.76 x 10^-2^), and Moderate (p = 4.21 x 10^-2^), while differences among lower grades were not significant (see Supplementary Table 2).

### 4.2 Comparison to the ground truth

For both the deconvolution and red-green methods, strong correlations were observed with ground truth fibrosis estimates, both before and after structure removal. The Spearman’s correlation coefficients were very strong for the deconvolution method (ρ = 0.78, p= 3.21 x 10^−4^) and moderately strong for the red-green method (ρ = 0.70, p = 2.29 x 10^−3^) before structure removal. After structural removal, the correlation with the ground truth improved for both the deconvolution (ρ = 0.81, p = 1.25 x 10^−4^) and the red-green (ρ = 0.81, p = 21.69 x 10^−4^) methods, with the red-green method showing a larger increase in correlation. Both methods tended to highly overestimate fibrosis prior to structure removal, as seen in Figure 4A and 4C. Removing non-interstitial structures decreased the mean difference results for both methods, with stain deconvolution achieving a narrower limit of agreement, as illustrated in the Bland-Altman plots in Figure 4B and 4D.

**Figure 4.**
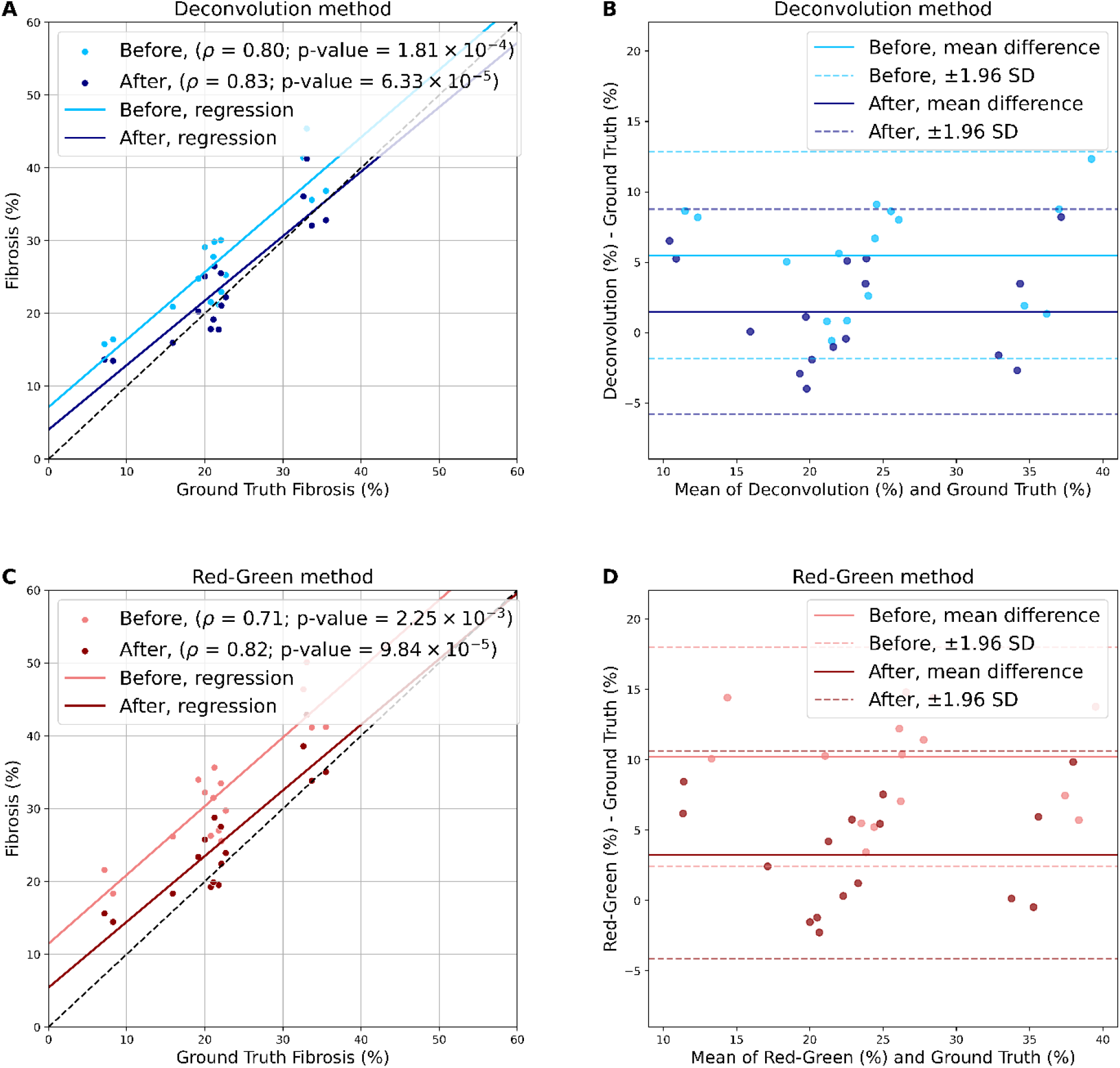
Correlation and agreement between digital fibrosis quantification methods and point counting reference. (A–B) The deconvolution method. (A) Regression plots comparing fibrosis percentages obtained by the deconvolution method with the ground truth (point counting), before (light blue) and after (dark blue) removal of non-interstitial structures. Spearman’s correlation coefficients (ρ) and corresponding p-values are shown. (B) Bland-Altman plots illustrating the agreement between the deconvolution method and ground truth measurements. Mean difference (solid line) and limits of agreement (±1.96 SD, dashed lines) are plotted separately for before and after structural removal. (C– D) Red-green method. (C) Regression plots as in (A), showing red-green method fibrosis estimates before (light red) and after (dark red) structural removal. (D) Bland-Altman plots as in (B), for the red-green method.

### 4.3. Comparison with serum creatinine levels

Serum creatinine is a widely used biomarker for assessing kidney function, with elevated levels typically indicating reduced GFR, as seen in both acute conditions such as tubular necrosis or interstitial nephritis, and in progressive kidney disease.

Across the investigated cases, fibrosis estimates were moderate-positively correlated with serum creatinine, supporting the algorithms’ biological plausibility. Figure 5A illustrates that for the deconvolution method, the Spearman’s correlation coefficient slightly improved from ρ = 0.56 (p = 0.02) before structure removal to ρ = 0.59 (p = 0.02) after removal. For the red-green method, (Figure 5B) structure removal was associated with a minimal change in correlation, from ρ = 0.60 (p = 0.01) to ρ = 0.59 (p = 0.02).

**Figure 5.**
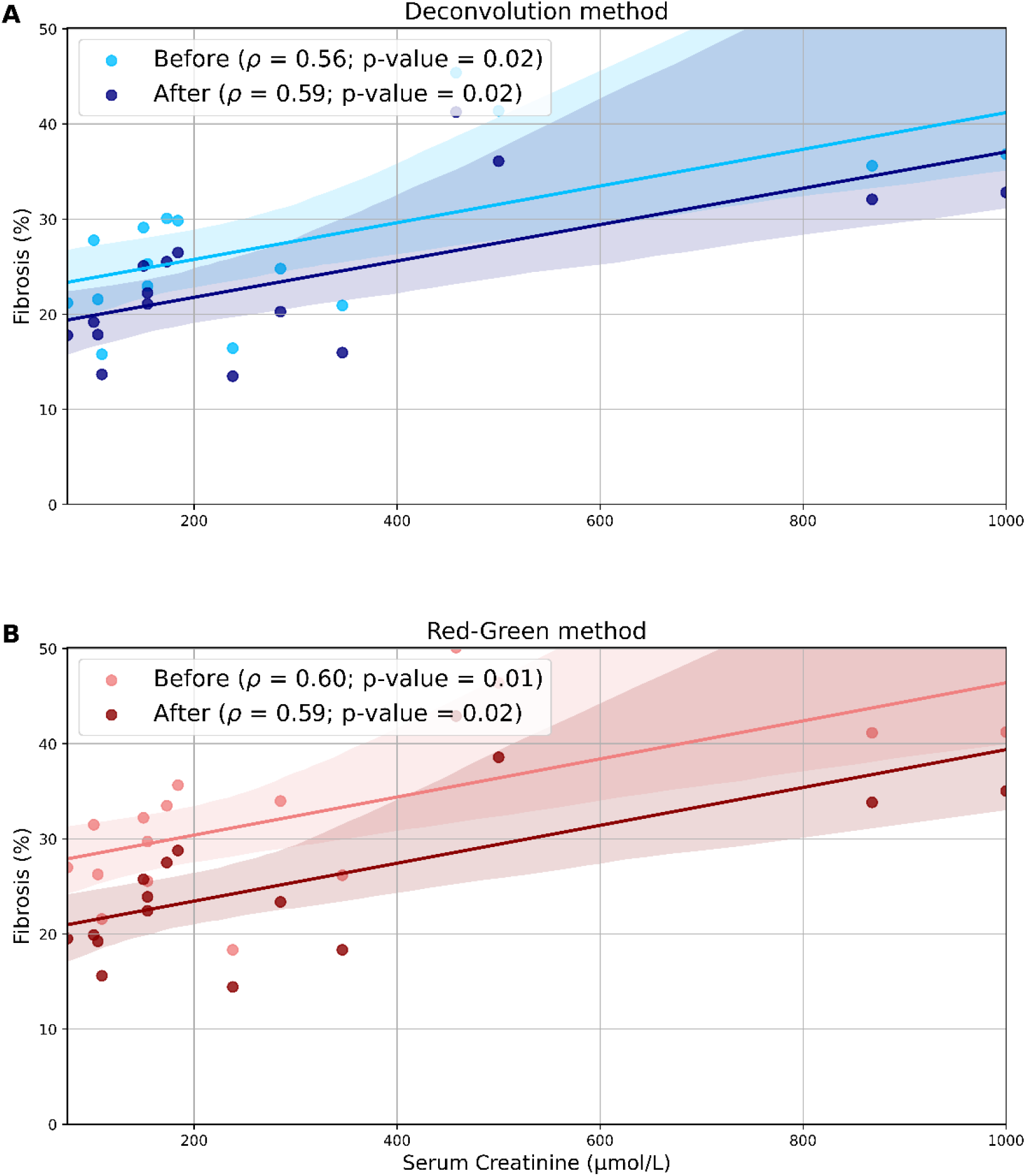
Correlation between digital fibrosis quantification and serum creatinine levels. (A–B) Scatter plots showing fibrosis percentages derived from the (A) deconvolution and (B) red-green methods, before and after structural segmentation, plotted against serum creatinine levels for each sample. Spearman’s correlation coefficients (ρ) and p-values are provided for each condition. Both methods show a moderate positive correlation with serum creatinine, with slightly improved correlation after removal of non-interstitial structures.

## 5. Discussion

Interstitial fibrosis (IF) is a key histopathological feature with proven relevance for estimating irreversible damage and predicting CKD prognosis. However, automatic quantification of IF remains challenging due to a lack of specificity of histological stains for interstitial structures. Sirius Red, for instance, often stains non-interstitial components in brightfield microscopy, including glomeruli, tubular epithelial cell nuclei and perivascular collagen. Ideally, segmenting non-interstitial structures prior to interstitial fibrosis measurement could provide a more accurate assessment of the extent of interstitial fibrosis. Although a few computer-aided tools^34,35^ have been proposed as more objective alternatives to traditional visual estimation, reliably restricting the analysis to the interstitial space has proven challenging. Despite its potential, this approach has not yet been extensively investigated.

In the present study, we examined the effect of excluding non-interstitial structures prior to fibrosis quantification with respect to two different quantification methods and two pathologist-derived fibrosis measurements. Our findings demonstrate that excluding non-interstitial tissue consistently and significantly improves the agreement between automatic IF quantification and expert assessments. We believe these findings substantially advance existing knowledge in the field by providing a more systematic evaluation of structural segmentation in fibrosis quantification compared to previously published studies.

### 5.1. Comparison to previous studies

Particularly, already in 2003, Grimm et al.^30^ proposed an automated image analysis pipeline for assessing IF in Sirius Red-stained kidney biopsies capable of predicting graft failure better than Banff scores. Their approach involved manually avoiding glomeruli, large vessels, subcapsular cortex, and medulla at the time of digitization. More recently, both Courtoy et al. (2020)^29^ and Dao et al. (2020)^28^ proposed well-performing methodologies for quantifying collagen proportionate area in Sirus red stained biopsies, also excluding some of the major confounding anatomical structures. Despite their contributions to the common goal of automatically quantifying interstitial fibrosis, these studies share several limitations: (i) reliance on non-objective visual estimation scores as a performance metric for a ground truth reference (ii) lack of thorough evaluation of performance gains following structural segmentation, and (iii) incomplete segmentation of non-interstitial structures. In contrast, the present study used manual point-counting for IF assessment, an established morphometric technique for area estimation^36^, and conducted a systematic comparison of automatic assessment performances before and after removal of non-interstitial structures. Finally, all non-interstitial components were fully removed, allowing for a more precise evaluation of fibrosis within the true region of interest: the renal interstitium.

### 5.2. Why and how our method work

As a consequence of the more systematic comparison conducted in the present study, we could demonstrate that segmentation of non-interstitial structures significantly reduced the systematic over-estimation of IF, thereby increasing performance. Thus, the study confirms that, without structural segmentation, IF quantification results might be negatively influenced by false-positive fibrotic signals likely originating from glomeruli, specifically when globally sclerosed, and various tubular structures.

At last, the current study also included an analysis of the relationship between interstitial fibrosis and serum creatinine. Both image analysis methods showed a moderate correlation with serum creatinine, which is in line with previously published findings^15,25^. While a stronger correlation might be expected given the role of fibrosis in kidney function decline, it is important to note that serum creatinine is influenced by multiple factors and is not a fibrosis-specific biomarker. The moderate correlation observed may reflect the heterogeneous nature of the dataset, which includes cases with additional pathological features such as interstitial edema, inflammation, and severe glomerular changes. These factors likely contribute to kidney dysfunction independently of fibrosis, thereby attenuating the strength of the observed associations. Thus, while the correlation supports biological plausibility, it does not necessarily serve as a direct measure of fibrosis assessment accuracy.

### 5.3. Limitations

The findings of the current study are subject to several limitations. For instance, the ground truth was based on manual point counting, an extremely time-consuming step that limited the size of the dataset. Due to similar time constraints, manual IF measurements in the sixteen WSIs were conducted by only a single expert pathologist, restricting investigations into inter- and intra-observer variability. While the available data were sufficient for a proof-of-concept comparison between automatic and manual assessments, they did not allow for a rigorous comparison of advantages and disadvantages of the two image analysis methods.

Additionally, tissue structures were segmented either completely manually (cortex, blood vessels, and glomeruli) or using a semi-automatic method (tubules). However, this manual effort is clearly impractical when considering an eventual clinical adoption of these tools. In recent years, a few deep-learning models have emerged as solutions capable of automatically segmenting, to a certain degree, various structures within the kidney’s cortex compartment^37^. Still, most of these models are tailored for more commonly used histological stains that display a variety of morphological changes, particularly Periodic Acid-Schiff (PAS). In contrast, Sirius Red has a very specific use-case: it is used solely for the purpose of fibrosis assessment, mostly for research applications. As a result, Sirius Red is not widely adapted in routine pathology workflows, and robust, publicly available tools for automatic segmentation remain scarce. Developing such tools would require substantial effort beyond the scope of the present study.

Finally, the accuracy of our workflow and the preliminary segmentation model may have been influenced by pre-analytical variables such as sample thickness, staining intensity, and tissue processing methods, as well as artifacts such as folds, tears, or bubbles. Given the small dataset, it was not feasible to investigate the effect of such variations on fibrosis quantification. A more comprehensive dataset incorporating such variability would be necessary to better understand these effects and to develop normalization strategies for downstream applications. Nevertheless, despite these limitations, both pipelines – though at an early proof-of-concept stage - show considerable potential for broader applications. Incorporating a post-processing step to detect edema—based on the presence of white pixels within the segmented interstitial area—could enable more nuanced investigations into tissue damage reversibility and therapeutic response. Furthermore, the same framework could be adapted to quantify fibrosis in other organs, such as the liver or lung, where collagen-based staining and morphometric analysis play similarly important diagnostic roles.

### 5.4. Overall conclusion and future directions

Taken together, our study presents accurate pipelines for automatic fibrosis quantifications in Sirius Red–stained kidney biopsies. We explicitly demonstrate that excluding non-interstitial structures through targeted segmentation significantly improves measurement accuracy, thus confirming the benefit of structure segmentation. While our pipeline showed robust performance in research settings, it is not yet intended for routine clinical application. Future work should validate this methodology in larger cohorts, integrating independent ground truth measures such as biochemical collagen assays, and evaluate its prognostic value across long-term renal outcomes. Additionally, extending these investigations to other routine stains – particularly trichrome – could broaden the applicability of the framework. Together, these efforts may pave the way toward reproducible, clinically applicable workflows for automated fibrosis assessment.

## 6. Budget

The project is funded by a PhD scholarship for Nazanin Mola (Helse Vest) and the strategic regional project “Pathology services in the Western Norway Health Region – a centre for applied digitization” (F-12563).

## 7. Ethical considerations

The study is approved by the Regional Ethical Committees (517496).

## 8. Acknowledgements

The results here are partially based upon machine-learning models trained on data generated by the Kidney Precision Medicine Project (https://www.kpmp.org).

