## Supplementary materials for "Image Analysis for Non-Neoplastic Kidney Disease: Utilizing Morphological Segmentation to Improve Quantification of Interstitial Fibrosis"

Supplementary Table 1. showing the results of the grid point counting in the manual evaluation of interstitial fibrosis in the16 cases. Samples are ordered by pathology grade, and within each grade, by ground truth fibrosis percentage.

| Cases | Creatinine values | Number of points interstitium white areas | Number of points interstitium fibrosis | Number of points others  (tubules + glomeruli) | Number of points in cortex | % Interstitium white areas | % Fibrosis | % Others | Age | IF | Gender | Cortical area |
| --- | --- | --- | --- | --- | --- | --- | --- | --- | --- | --- | --- | --- |
| 1 | 109 | 86 | 37 | 394 | 517 | 16.63 | 7.16 | 76.21 | 17 | 0 | m | 9.92 |
| 2 | 238 | 116 | 42 | 351 | 509 | 22.79 | 8.25 | 68.96 | 38 | 0 | m | 8.04 |
| 3 | 345 | 104 | 80 | 319 | 503 | 20.68 | 15.9 | 63.42 | 31 | 0 | m | 0.97 |
| 4 | 75 | 55 | 83 | 243 | 381 | 14.44 | 21.78 | 63.78 | 50 | 0 | k | 5.42 |
| 5 | 105 | 31 | 119 | 423 | 573 | 5.41 | 20.77 | 73.82 | 48 | 1 | m | 9.05 |
| 6 | 101 | 57 | 100 | 317 | 474 | 12.03 | 21.1 | 66.88 | 53 | 1 | m | 10.45 |
| 7 | 154 | 64 | 132 | 386 | 582 | 11 | 22.68 | 66.32 | 33 | 1 | m | 8.46 |
| 8 | 868 | 56 | 157 | 253 | 466 | 12.02 | 33.69 | 54.29 | 61 | 1 | k | 3.54 |
| 9 | 285 | 94 | 101 | 332 | 527 | 17.84 | 19.17 | 63 | 30 | 2 | m | 5.11 |
| 10 | 150 | 99 | 112 | 349 | 560 | 17.68 | 20 | 62.32 | 51 | 2 | m | 7.44 |
| 11 | 184 | 143 | 121 | 306 | 570 | 25.09 | 21.23 | 53.68 | 56 | 2 | m | 4.5 |
| 12 | 154 | 73 | 105 | 297 | 475 | 15.37 | 22.11 | 62.53 | 33 | 2 | m | 2.56 |
| 13 | 173 | 119 | 103 | 245 | 467 | 25.48 | 22.06 | 52.46 | 71 | 3 | k | 5.88 |
| 14 | 500 | 165 | 182 | 211 | 558 | 29.57 | 32.62 | 37.81 | 51 | 3 | m | 3.16 |
| 15 | 458 | 149 | 192 | 240 | 581 | 25.65 | 33.05 | 41.31 | 70 | 3 | k | 4.52 |
| 16 | 1000 | 141 | 180 | 186 | 507 | 27.81 | 35.5 | 36.69 | 21 | 3 | m | 4.79 |

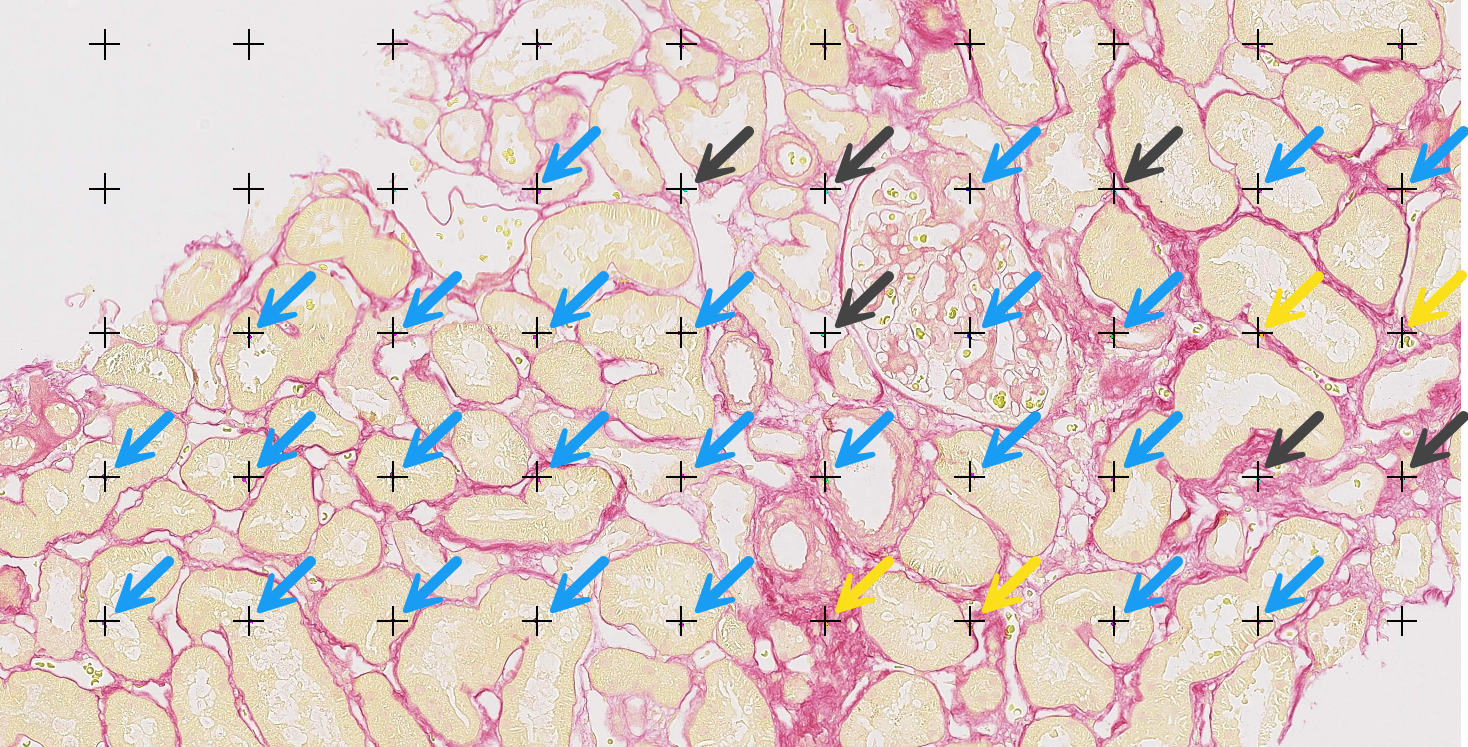

Supplementary Figure 1: **Point counting on the Sirius red stained kidney biopsy.** The blue color arrows mark the “others” category (including tubules and glomeruli). The yellow color arrows mark connective tissue. The black arrows mark the Edema (white areas) in the interstitium. The sum of yellow and black arrows represents the interstitial area. The grid points that didn’t hit the tissue were left unassigned.

Supplementary Table 2. Post hoc pairwise comparisons to see how (A) deconvolution (B) and red-green methods estimated fibrosis across different pathology grades, using Tukey’s Honestly Significant Difference test. The table shows the mean difference between pathology grade groups, adjusted p-values, confidence intervals, and whether the difference was statistically significant at α = 0.05 (reject = True). Significant differences were found between the Severe group and one or more of the other pathology grades for both methods.

A

| Group 1 | Group2 | Mean diff | p-adj | lower | upper | Reject |
| --- | --- | --- | --- | --- | --- | --- |
| Mild | Minimal | -7.605 | 0.186 | -17.9963 | 2.7863 | False |
| Mild | Moderate | 0.3975 | 0.9994 | -9.9938 | 10.7888 | False |
| Mild | Severe | 11.0825 | 0.0355 | 0.6912 | 21.4738 | True |
| Minimal | Moderate | 8.0025 | 0.1559 | -2.3888 | 18.3938 | False |
| Minimal | Severe | 18.6875 | 0.0009 | 8.2962 | 29.0788 | True |
| Moderate | Severe | 10.685 | 0.0432 | 0.2937 | 21.0763 | True |

B

| Group 1 | Group2 | Mean diff | p-adj | lower | upper | Reject |
| --- | --- | --- | --- | --- | --- | --- |
| Mild | Minimal | -7.2475 | 0.2285 | -17.8128 | 3.3178 | False |
| Mild | Moderate | 0.8625 | 0.9947 | -9.7028 | 11.4278 | False |
| Mild | Severe | 11.7825 | 0.0276 | 1.2172 | 22.3478 | True |
| Minimal | Moderate | 8.11 | 0.1577 | -2.4553 | 18.6753 | False |
| Minimal | Severe | 19.03 | 0.0009 | 8.4647 | 29.5953 | True |
| Moderate | Severe | 10.92 | 0.0421 | 0.3547 | 21.4853 | True |

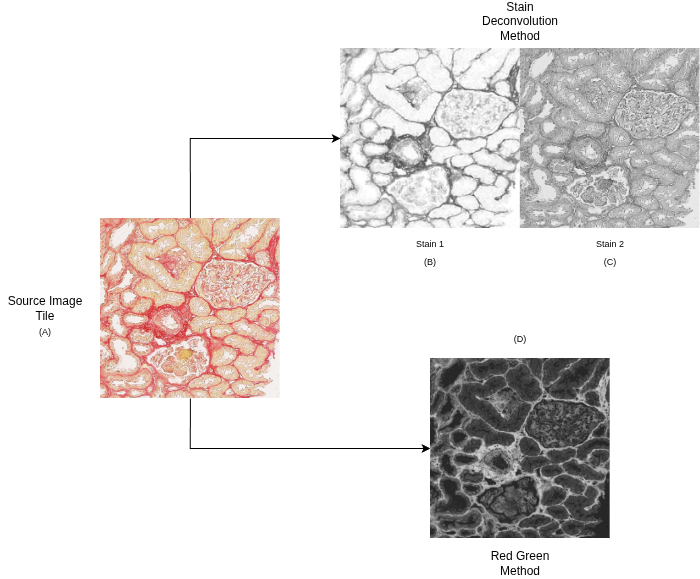

*Supplementary Figure 2:* ***The output of the two image analysis methods****. First row shows the output of the stain deconvolution method: (A) Original tile extracted from one of the WSIs; (B) First stain component (Sirius Red) and (C) Second stain component (Picric acid). The second row shows the output of red-green method and the: (D) Result of the matrix subtraction between the red and green channels, showing improved contrast for collagen-stained areas.*
